# Parental effects of pathogenic bacterial infections in *Drosophila melanogaster*: trans-generational immune priming without apparent costs

**DOI:** 10.1101/2024.02.07.579292

**Authors:** Aabeer Kumar Basu, Nagaraj Guru Prasad

## Abstract

Parental experience with pathogens and parasites can shape offspring susceptibility to infections, a phenomenon commonly referred to as trans-generational immune priming (TgIP) in insects. TgIP can influence the evolution of both host immune traits and pathogen virulence. The host ecology and its environmental pathogen load can in turn shape the evolution of TgIP. Therefore, the functional limits of TgIP and the conditions under which it is observed need to be explored in a variety of species. Majority of empirical studies addressing this phenomenon focus on non-pathogenic infections of parents, implemented via either the use of pathogen substitutes (viz., liposaccharide molecules), or the use of sub-lethal infection doses, by using inactivated pathogens to infect the parents. We tested if exposing parents to pathogenic bacterial infections affects offspring susceptibility to homologous (same pathogen as the one used to infect the parents) and heterologous (different pathogen than the one used to infect the parents) infection challenges in *Drosophila melanogaster*. We found that parental infection increases offspring survival following a homologous challenge. Similar effects were seen in the case of heterologous challenges too, with the breadth of cross-reactivity being dependent on the pathogen used to infect the parents. Parental infection also improves offspring capacity to restrict systemic pathogen proliferation following a homologous challenge, suggesting increased disease resistance. Parental infection has no effect on offspring reproduction, either in presence or absence of infection, suggesting an absence of associated costs and limited benefit of trans-generational immune priming. Overall, we have demonstrated that parental infection helps offspring better counter infection challenges in *D. melanogaster* without incurring any measurable costs.

## 1. Introduction

Parental effects, that is plastic changes in offspring phenotypes by virtue of the environment experienced by the parents, are often beneficial to the offspring and helps the offspring maintain fitness under stressful conditions in various organisms (Yin et al., 2019). These plastic improvements in offspring fitness can influence long term organismal evolution by a variety of mechanisms (Bonduriansky and Day 2009). Parental effects can influence a wide range of organismal traits, including defense against pathogens and parasites, in both vertebrates and invertebrates (Roth et al., 2018). Improvement of offspring defense against pathogens and parasites, by virtue of parental experience with pathogens and other immunogenic entities, is commonly referred to as trans-generational immune priming in the invertebrates (Little and Kraaijeveld 2004, Kurtz 2005). Over the years various empirical studies have explored this phenomenon in insects and other invertebrates, and various associated nuances have been elucidated (Milutinovic et al., 2016, Contreras-Garduno et al., 2016, Cooper and Eleftherianos 2017, Roth et al., 2018, Tetreau et al., 2019, Prakash and Khan 2022, Rutkowski et al., 2023). Theoretical studies have also demonstrated that immune priming across generations can have epidemiological consequences and impact the dynamics of both the host and the pathogen populations (Tate and Rudolf 2011, Tidbury et al., 2012, Pigeault et al., 2016). Yet, various aspects of trans-generational immune priming continue to be debated. For example, is priming specific (viz., Dhinaut et al., 2018a, Dhinaut et al., 2018b, Ben-Ami et al., 2020)? Does it have costs (viz., Zanchi et al., 2012, Dhinaut et al., 2018a, Dhinaut et al., 2018b)? Does priming improve offspring disease resistance or tolerance (viz., Paraskevopoulou et al., 2022)?

It is common practice in empirical investigations of trans-generational immune priming to infect the hosts of the parental generation with either substances which mimic pathogens, or inactivated pathogens, or with a non-lethal dose of an active pathogen (Tetreau et al., 2019, Rutkowski et al., 2023). Such an experimental design has its benefits. *One*, it separates the confounding effects of the host’s response to infection and the pathogen’s modification of the host physiology. *Two*, it ensures that improvement in offspring defenses is not due to survivor bias and selection for improved defenses in the parental generation. While these are important factors, this experimental design overlooks important and inevitable consequences associated with natural infections: pathogenicity, sickness, and lethality. Although a recent study has demonstrated in beetles that lethal infection in parents does not lead to improvement in offspring post-infection survival when infected with the same pathogen (Schulz et al., 2023), we were interested to test if subjecting *Drosophila melanogaster* parents to infection with known natural bacterial pathogens had an effect on offspring disease susceptibility. Previous studies have suggested that subjecting female parents to bacterial infections does not lead to trans-generation immune priming in this species (Linder and Promislow 2009, Radhika and Lazzaro 2023), while viral infections do induce priming across generations (Mondotte et al., 2020). For our study, we infected *D. melanogaster* parents with two bacterial pathogens, *Erwinia carotovora carotovora* and *Enterococcus faecalis*, and tested if doing so alters offspring post-infection survival, disease resistance, and reproductive capacity. We find that parental infection reduces offspring susceptibility to infections and increases offspring disease resistance, without any measurable impact on offspring reproductive output.

## 2. Materials and methods

### 2.1. Host population and general handling

Flies from the BRB2 population - a large, lab adapted, out-bread *Drosophila melanogaste*r population - were used for the experiments. The Blue Ridge Baseline 2 (BRB2) population were originally established by hybridizing 19 wild-caught iso-female lines (Singh et al., 2015), and has been maintained since as an outbred population on a 14-day discrete generation cycle with census size of about 2800 adults per generation. Every generation, eggs are collected from population cages (plexiglass cages: 25 cm length × 20 cm width × 15 cm height) and dispensed into vials (25 mm diameter × 90 mm height) with 8 mL standard banana-jaggery-yeast food medium, at a density of 70 eggs per vial. 40 such vials are set up; the day of egg collection is demarcated as day 1. The vials are incubated at 25 °C, 50-60% RH, 12:12 hour LD cycle; under these conditions the egg-to-adult development time for these flies is about 9-10 days. On day 12 post egg-collection all adults are transferred to the population cage, and provided with fresh food plates (banana-jaggery-yeast food medium in a 90 mm Petri plate) supplemented with *ad libitum* live yeast paste. On day 14, the cage is provided with a fresh food plate, and 18 hours later eggs are collected from this plate to initiate the next generation.

### 2.2. Pathogen handling and infection protocol

Four bacterial pathogens were used in this study for infecting the flies:

a. *Enterococcus faecalis* (Lazzaro et al., 2006), incubation temperature 37 ^O^C;
b. *Erwinia carotovora carotovora*, strain Ecc15 (Martins et al., 2013), incubation temperature 29 ^O^C;
c. *Bacillus thuringiensis* (DSMZ, Germany, catalogue number: DSM2046), incubation temperature 30 ^O^C; and,
d. *Pseudomonas entomophila*, strain L48 (Vodovar et al., 2005, Mullet et al., 2012), incubation temperature 27 ^O^C.

All pathogens are maintained in the lab as glycerol stocks, and are cultured in lysogeny broth (Luria Bertani Broth, Miler, HiMedia M1245). Overnight culture of bacteria grown from glycerol stocks was diluted (1:100) in fresh LB medium and incubated till confluency (optical density OD_600_ = 1.0-1.2). The bacterial cells were pelleted down by centrifugation and re-suspended in sterile 10 mM MgSO_4_ buffer at OD_600_ = 1.0. OD600 = 1.0 corresponds to 10^7^ cells/mL for *E. faecalis*, 10^7^ cells/mL for *E. c. carotovora*, 10^4^ cells/mL for *B. thuringiensis*, and 10^6^ cells/mL for *P. entomophila*. Flies were infected by pricking them at the dorsolateral side of the thorax with a fine needle (Minutien pin, 0.1 mm, Fine Science Tools, CA, item no. 26001-10) dipped in bacterial suspension under light CO_2_ anesthesia. Flies for sham-infections were similarly treated but pricked with needle dipped in sterile 10 mM MgSO_4_ buffer.

### 2.3. Derivation and handling of parental (P) generation flies

Eggs were collected from BRB2 population cages at a density of 70 eggs per vial with 8 mL of standard food medium, and reared till adulthood identically to how the BRB2 population is maintained in the lab. Eggs developed into adults in 9-10 days following egg collection and were housed in the rearing vial till day 12 for them to become mature adults. On day 12 after egg collection, 2–3-day old adults were randomly distributed into two treatments: (i) infected with pathogen (n = 200 females and 200 males), and (ii) sham-infected (n = 100 females and 100 males); these two treatments represent the ‘infected parent’ and ‘control parent’ treatments, respectively. Thereafter, these adults were housed in treatment-specific plexiglass cages (14 cm length × 16 cm width × 13 cm height) with *ad libitum* access to standard food medium (in 60 mm Petri plates). The number of flies in the infected treatment were greater than in the sham-infected treatment to account for the fact that about fifty to sixty percent of the infected flies die from infection, and therefore sufficient numbers of flies must remain alive during the egg-collection window, which was after the acute mortality phase, to contribute to the next generation (**supplementary figure S1**). Flies in the ‘infected parent’ treatment were either infected with *Erwinia c. carotovora* or with *Enterococcus faecalis* depending on the experimental requirement. For experiments in which the flies were infected with *E. c. carotovora*, eggs for the next generation were collected 96 hours post-infection from both parental treatments. For experiments in which the flies were infected with *E. faecalis*, eggs for the next generation were collected either 48 hours (for all offspring trait assays) or 96 hours (only for survival after homologous infection challenge) post-infection from both parental treatments, as per the experimental needs. Eggs for offspring generations were also collected at a density of 70 eggs per vial with 8 mL of standard food medium, and reared till adulthood identically to how the BRB2 population is maintained in the lab. Eggs developed into adults in 9-10 days following egg collection and were housed in their rearing vial till day 12 for them to become mature adults.

### 2.4. Post-infection survival of first offspring (F1) generation following homologous infection challenge

2–3-day old adult offspring from each parental treatment (infected vs. control) were randomly assigned to two treatments: (i) infected (n = 200 females and 200 males), and (ii) sham-infected (n = 100 females and 100 males). Offspring of parents infected with *E. c. carotovora*, and their corresponding control parents, were infected with *E. c. carotovora*, and offspring of parents infected with *E. faecalis*, and their corresponding control parents, were infected with *E. faecalis*. Thereafter, these adults were housed in treatment-specific plexiglass cages with *ad libitum* access to standard food medium, and their mortality was recorded every 4-6 hours till 120 hours post-infection. Fresh food was provided to the cages every 48 hours. For each pathogen, starting from the parental generation, the experiment was independently replicated twice.

### 2.5. Post-infection survival of second offspring (F2) generation following homologous infection challenge

2–3-day old adult offspring from each grand-parental (generation P) treatment (infected vs. control) were randomly assigned to two treatments: (i) infected (n = 100 females and 100 males), and (ii) sham-infected (n = 50 females and 50 males). Offspring of grand-parents infected with *E. c. carotovora*, and their corresponding control grand-parents, were infected with *E. c. carotovora*, and offspring of grand-parents infected with *E. faecalis*, and their corresponding control grand-parents, were infected with *E. faecalis*. Thereafter, these adults were housed in treatment-specific plexiglass cages with *ad libitum* access to standard food medium, and their mortality was recorded every 4-6 hours till 120 hours post-infection. Fresh food was provided to the cages every 48 hours. For each pathogen, starting from the parental generation, the experiment was independently replicated twice.

### 2.6. Post-infection survival of first offspring (F1) generation following heterologous infection challenge

2–3-day old adult offspring from each parental treatment (infected vs. control) were either infected with one of the three novel pathogens (pathogen not used for infection in generation P; n = 60 females and 60 males for each pathogen) or subjected to sham infections (n = 60 females and 60 males). Offspring of parents infected with *E. c. carotovora*, and their corresponding control parents, were infected with *B. thuringiensis*, *E. faecalis*, and *P. entomophila*. Offspring of parents infected with *E. faecalis*, and their corresponding control parents, were infected with *B. thuringiensis*, *E. c. carotovora*, and *P. entomophila*. Thereafter, these adults were housed in treatment-specific plexiglass cages with *ad libitum* access to standard food medium, and their mortality was recorded every 4-6 hours till 120 hours post-infection. Fresh food was provided to the cages every 48 hours. For each pathogen, starting from the parental generation, the experiment was independently replicated twice.

### 2.7. Early-life fecundity of first offspring (F1) generation females in absence of infection

2–3-day old flies were sorted into vials at a density of 5 females and 5 males per vial, with 8 mL of standard food medium in each vial; 10 such vials were set up for offspring of each parental treatment (infected vs. control). Flies were housed in these vials for 24 hours, during which the females laid eggs, and then transferred to fresh food vials, with one-to-one mapping of vial identity. This procedure was repeated for 8 consecutive days. The vials with eggs were maintained under the regular rearing environment of the BRB2 flies. 12 days after egg laying, the number of adult progenies in each vial was scored as a measure of reproductive output of females in that vial. The total number of progenies in the vials was divided by the number of alive females (usually 5, unless any female was lost during the transfer process) in the vial to obtain the measure of per capita female fecundity for each day. For each pathogen, starting from the parental generation, the experiment was independently replicated twice.

### 2.8. Competitive fertilization success of first offspring (F1) generation males in absence of infection

Male offspring from each parental treatment, infected vs. control, were individually housed in standard food vials (n = 50 males per parental treatment), along with a BL*st* male and a pre-inseminated BL*st* female. (BL*st* is an outbred population, maintained under identical maintenance conditions as BRB2. The BL*st* population is fully homozygous for the recessive eye-colour marker, *scarlet*.) The BL*st* flies were age matched with the focal males. The vials were left undisturbed for 24 hours, so that the two males can compete for the opportunity to inseminate the female and fertilize the eggs produced by it. 24 hours later both males were discarded, and the female was housed individually in a fresh food vial (with 8 mL of standard food medium) and allowed to lay eggs for 24 hours, following which the female was discarded. The vials with eggs were maintained under the regular rearing environment of the BRB2 flies. 12 days after egg laying, the adult progenies were scored on the basis of eye colour to determine paternity. The proportion of progenies sired by the focal male was considered as a measure of its competitive fertilization success. For each pathogen, starting from the parental generation, the experiment was independently replicated twice.

### 2.9. Post-infection systemic pathogen growth in first offspring (F1) generation flies following homologous infection challenge

This experiment was carried out only with offspring of parents infected with *E. faecalis*, and their corresponding control parents, and the experiment was independently replicated four times, starting from the parental generation. 4–5-day old adult flies, from both parental treatments, were either subjected to infections with *E. faecalis* (n = 30 females and 30 males) or were subjected to sham-infections (n = 30 females and 30 males). Thereafter, these adults were housed in treatment-specific plexiglass cages with *ad libitum* access to standard food medium. At 4- and 10-hours post-infection, 12 infected females and 12 infected males from each parental treatment were randomly sampled and their systemic bacterial load was measured. Systemic bacterial load for 12 sham-infected females and 12 sham-infected males from each parental treatment was also measured at both time-points.

For measuring systemic bacterial loads, flies were washed twice in 70% ethanol for 1 minute and then transferred individually to 1.5 mL microcentrifuge tubes containing 50 microliters of sterile MgSO_4_ (10 mM) solution. Flies were homogenized in these vials using a motorized pestle for 60 seconds. This homogenate was serially diluted (1:10 dilutions) 8 times in sterile MgSO_4_ (10 mM) solution; 10 microliters in 100 microliters. 10 microliters from each dilution, and the original homogenate, were spotted onto a lysogeny agar plate (2% agar, Luria Bertani Broth, Miler, HiMedia). The plates were incubated at 37 ^O^C, for 8-12 hours, and the number of colony-forming units (CFUs) in each dilution was counted. The number of CFUs in the countable dilution (30 ≤ CFUs ≥ 300) was multiplied by appropriate dilution factor to obtain the bacterial load for each individual fly.

### 2.10. Fecundity of first offspring (F1) generation females following homologous infection challenge

This experiment was carried out only with offspring of parents infected with *E. faecalis*, and their corresponding control parents, and the experiment was independently replicated four times, starting from the parental generation. 4-5-day old adult, inseminated females from both parental treatments were either subjected to infection with *E. faecalis* (n = 80 females) or were subjected to sham-infections (n = 40 females). Thereafter these females were housed individually in food vials (with 8 mL standard food medium), where they laid eggs for the next 48 hours, during which their mortality was also monitored, every 2 hours. At the end of this period, all the alive females were discarded, and the carcasses of the dead females were removed from their vials. The vials were then incubated under regular BRB2 population maintenance conditions, and 12 days later, the number of adult progeny in each vial was enumerated. Since survival of infected females varied considerably amongst themselves and from that of sham-infected females (for which no death occurred within the observation window), the number of progeny produced by each female was divided by the hours survived by that female to obtain the final measure of fecundity, which was subjected to statistical analysis.

### 2.11. Data analysis

All statistical analyses were carried out using R statistical software (version 4.1.0, R Core Team 2021). Post-infection survival data for F1 or F2 offspring was analyzed using mixed-effects Cox proportional hazards model, with ‘parental treatment’ as a fixed factor and ‘replicate’ as a random factor; data for female and male offspring were analyzed separately. Since in most experiments, there was negligible or zero mortality in sham-infected treatments (**supplementary figure S2**), data from only the infected offspring flies were subjected to survival analysis. Female early-life fecundity (progeny per female) was analyzed using mixed-effects, type-III Analysis of Variance (ANOVA) with ‘parental treatment’, ‘day of measurement’, and their interaction as fixed factors, and ‘replicate’ as a random factor. Male competitive fertilization success (proportion progeny sired) was analyzed using mixed-effects, type-III ANOVA with ‘parental treatment’ as a fixed factor, and ‘replicate’ as a random factor. Data for systemic bacterial load was analyzed using mixed-effects, type-III ANOVA with ‘parental treatment’, ‘hours post-infection’, and their interaction as fixed factors, and ‘replicate’ as random factor; data from female and male offspring were analyzed separately. Post-infection female fecundity (progeny per female per hour) was analyzed using mixed-effects, type-III ANOVA with ‘parental treatment’, ‘offspring infection status’, and their interaction as fixed factors, and ‘replicate’ as random factor.

## 3. Results

### 3.1. Post-infection survival of first (F1) and second (F2) offspring generation flies after homologous infection challenge

Flies of parental generation (generation P) were infected with *Erwinia c. carotovora* (*Ecc*), and their F1 and F2 offspring were tested for post-infection survival after being infected with the same pathogen. When infected with *Ecc*, F1 offspring of *Ecc*-infected parents were better at surviving the infection compared to offspring of control (sham-infected) parents, in case of both female (hazard ratio, 95% confidence interval: 0.630, 0.529-0.752) and male (HR, 95% CI: 0.534, 0.447-0.637) offspring (**figure 1.A**). Female F2 offspring of *Ecc*-infected parents were not different from the female F2 offspring of control parents in terms of post-infection survival (HR, 95% CI: 1.099, 0.876-1.380), while male F2 offspring of *Ecc*-infected parents were more susceptible to infection compared to male F2 offspring of control parents (HR, 95% CI: 1.284, 1.040-1.587) (**figure 1.B**).

**Figure 1.**
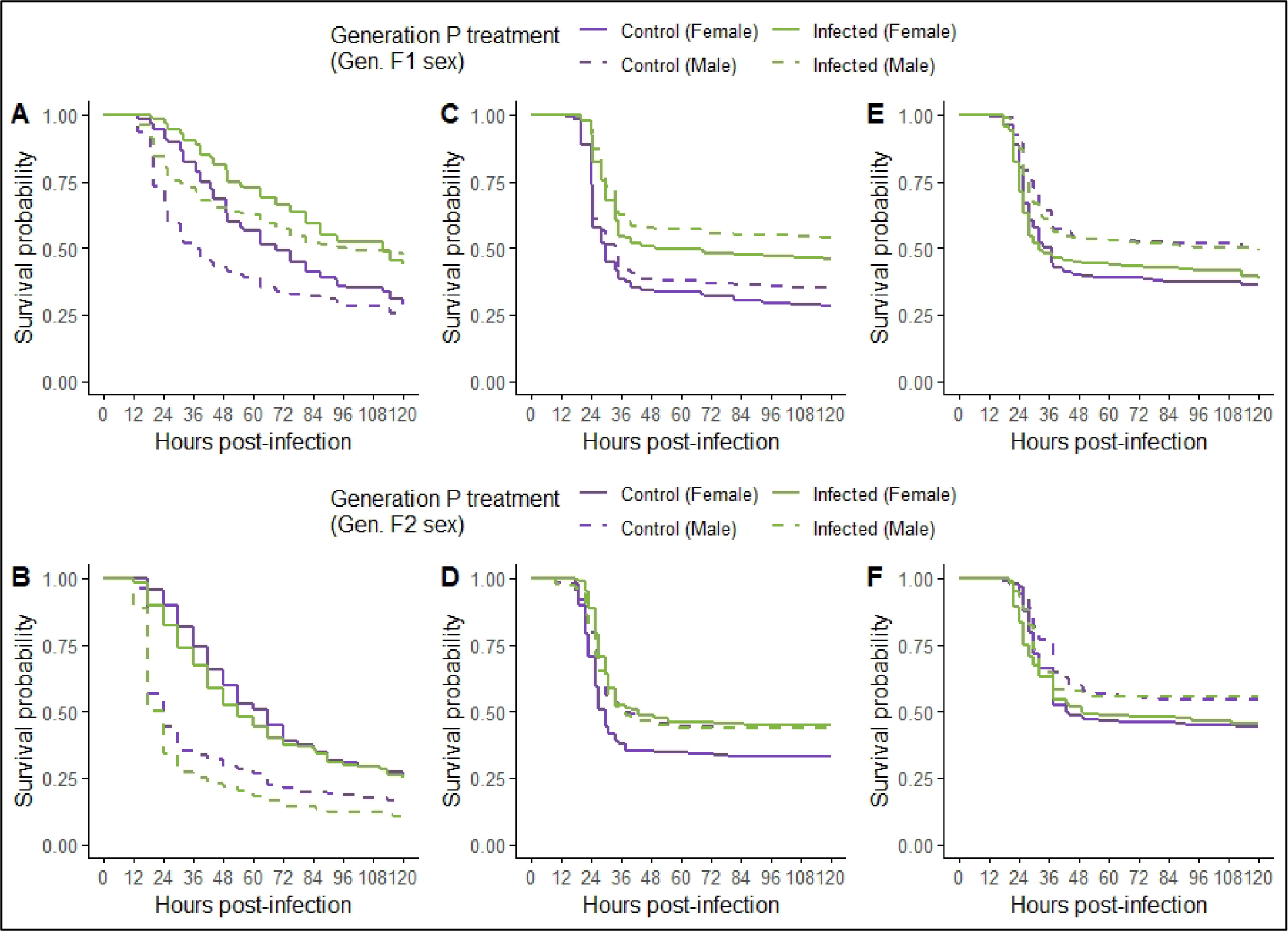
Post-infection survival of first- (F1) and second-offspring (F2) generation flies of infected and control parents after a homologous infection challenge. Offspring of *Erwinia c. carotovora* (*Ecc*) infected parents were infected with *E. c. carotovora*. Offspring of *Enterococcus faecalis* (*Ef*) infected parents were infected with *E. faecali*s. **(A)** F1 offspring of *Ecc*-infected parents. **(B)** F2 offspring of *Ecc*-infected parents. **(C)** F1 offspring of *Ef*-infected parents (eggs collected at 48 hours post-infection (HPI) in the parental generation). **(D)** F2 offspring of *Ef*-infected parents (eggs collected at 48 HPI in the parental generation). **(E)** F1 offspring of *Ef*-infected parents (eggs collected at 96 HPI in the parental generation). **(F)** F2 offspring of *Ef*-infected parents (eggs collected at 96 HPI in the parental generation). Survival curves plotted using Kaplan-Meier method after pooling data from both replicates for each pathogen.

Flies of parental generation (generation P) were infected with *Enterococcus faecalis* (*Ef*), and their F1 and F2 offspring were tested for post-infection survival after being infected with the same pathogen. In case of eggs collected 48 hours post-infection in generation P, when infected with *Ef*, F1 offspring of *Ef*-infected parents were better at surviving the infection compared to offspring of control parents, in case of both female (hazard ratio, 95% confidence interval: 0.563, 0.673-0.742) and male (HR, 95% CI: 0.535, 0.443-0.647) offspring (**figure 1.C**). Female F2 offspring of *Ef*-infected parents had better post-infection survival compared to female F2 offspring of control parents (HR, 95% CI: 0.634, 0.492-0.816), but male F2 offspring of *Ef*-infected and control parents did not differ from one another in terms of survival (HR, 95% CI: 1.026, 0.790-1.332) (**figure 1.D**).

In case of eggs collected 96 hours post-infection in generation P, when infected with *Ef*, F1 offspring of *Ef*-infected parents, both females (HR, 95% CI: 1.005, 0.844-1.198) and males (HR, 95% CI: 1.053, 0.866-1.281), did not differ in terms of post-infection survival compared to offspring of control parents (**figure 1.E**). Similarly, F2 offspring of *Ef*-infected parents, both females (HR, 95% CI: 1.042, 0.800-1.358) and males (HR, 95% CI: 1.067, 0.796-1.429), did not differ in terms of post-infection survival compared to offspring of control parents (**figure 1.F**).

### 3.2. Post-infection survival of first offspring (F1) generation flies after heterologous infection challenge

Flies of parental generation (generation P) were infected with *E. c. carotovora* (*Ecc*), and their F1 offspring were tested for post-infection survival after being infected with *Bacillus thuringiensis* (*Bt*) (**figure 2.A**), *E. faecalis* (*Ef*) (**figure 2.B**), and *Pseudomonas entomophila* (*Pe*) (**figure 2.C**). Female offspring of *Ecc*-infected parents were not significantly different in terms of post-infection survival from female offspring of control parents when infected with *Bt* (HR, 95% CI: 0.831, 0.626-1.103), *Ef* (HR, 95% CI: 0.887, 0.657-1.198), or *Pe* (HR, 95% CI: 1.132, 0.861-1.489). Male offspring of *Ecc*-infected parents survived significantly better compared to male offspring of control parents when infected with *Ef* (HR, 95% CI: 0.729, 0.535-0.993) and *Pe* (HR, 95% CI: 0.618, 0.474-0.805), but not when infected with *Bt* (HR, 95% CI: 0.864, 0.649-1.149).

**Figure 2.**
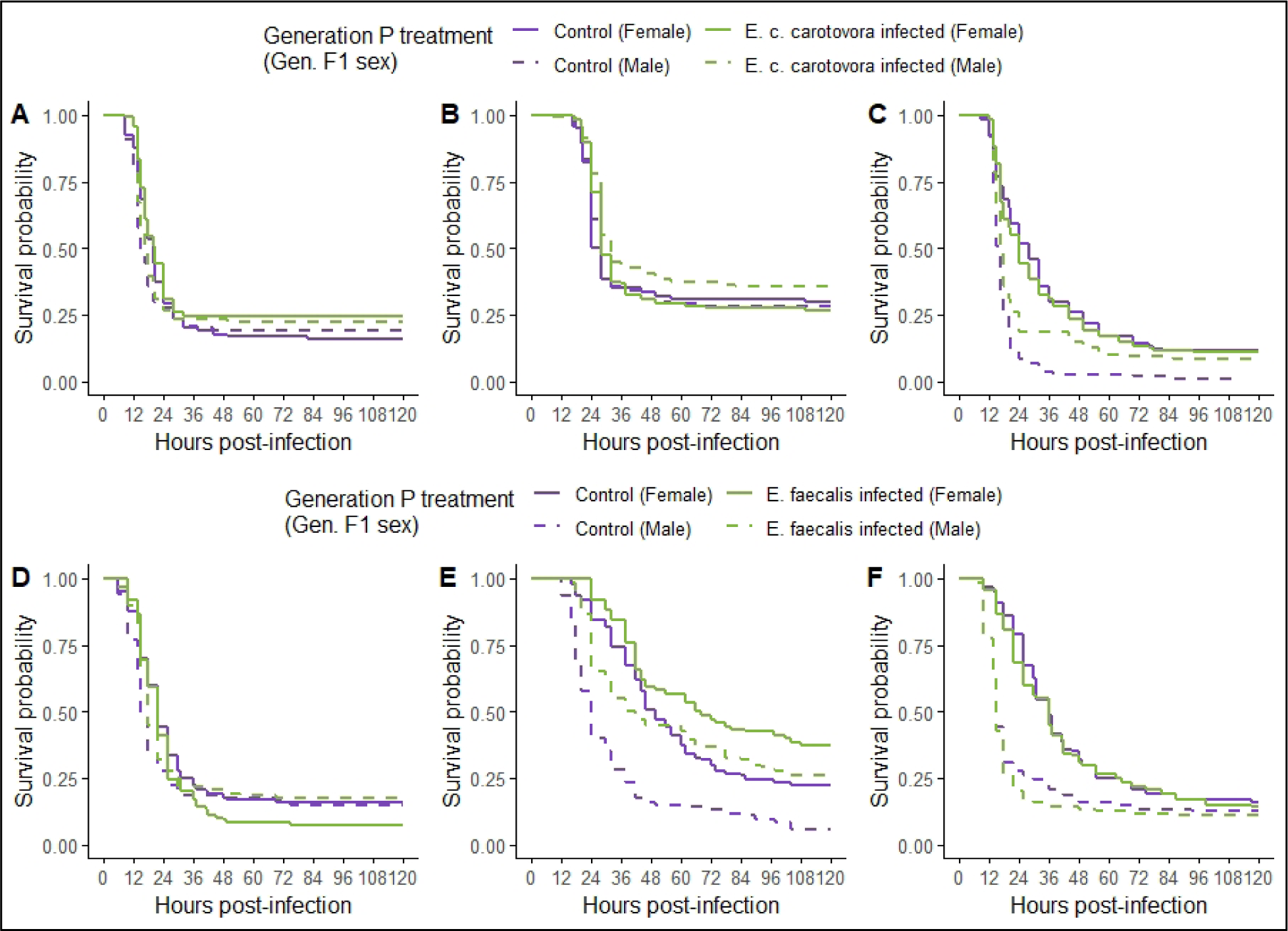
Post-infection survival of first-offspring (F1) generation flies of infected and control parents after heterologous infection challenges. Offspring of *Erwinia c. carotovora* infected parents were infected with **(A)** *Bacillus thuringiensis*, **(B)** *Enterococcus faecalis*, and **(C)** *Pseudomonas entomophila*. Offspring of *E. faecalis* infected parents were infected with **(D)** *B. thuringiensis*, **(E)** *E. c. carotovora*, and **(F)** *P. entomophila*. Survival curves plotted using Kaplan-Meier method after pooling data from both replicates for each pathogen.

Flies of parental generation (generation P) were infected with *E. faecalis* (*Ef*), and their F1 offspring were tested for post-infection survival after being infected with *B. thuringiensis* (*Bt*) (**figure 2.D**), *E. c. carotovora* (*Ecc*) (**figure 2.E**), and *P. entomophila* (*Pe*) (**figure 2.F**). Offspring of *Ef*-infected parents were not significantly different in terms of survival from offspring of control parents, when infected with *Bt* in case of both female (HR, 95% CI: 1.263, 0.962-1.658) and male (HR, 95% CI: 0.774, 0.587-1.021) offspring, and when infected with *Pe* in case of both female (HR, 95% CI: 1.105, 0.839-1.454) and male (HR, 95% CI: 1.258, 0.954-1.659) offspring. When infected with *Ecc*, offspring of *Ef*-infected parents survived significantly better compared to offspring of control parents, in case of both female (HR, 95% CI: 0.658, 0.485-0.892) and male (HR, 95% CI: 0.451, 0.340-0.597) offspring.

### 3.3. Reproductive output of first offspring (F1) generation flies in absence of infection

Flies of parental generation (generation P) were infected with *E. c. carotovora*, and the reproductive output of F1 offspring was measured: early-life fecundity for females and competitive fertilization success (CFS) for males. Day of fecundity measurement had a significant effect on per capita female fecundity (F_1,320_ = 392.2122, p-value < 2.2e-16), but parental infection treatment did not have a significant effect (F_1,320_ = 0.6158, p-value = 0.433), and neither did the interaction between age × parental treatment (F_1,320_ = 0.0004, p-value = 0.984) (**figure 3.A**). Parental infection treatment had a significant effect on male CFS (F_1,195_ = 4.242, p-value = 0.041), with male offspring of infected parents being better at competing for fertilizations compared to male offspring of sham-infected parents (**figure 3.B**).

**Figure 3.**
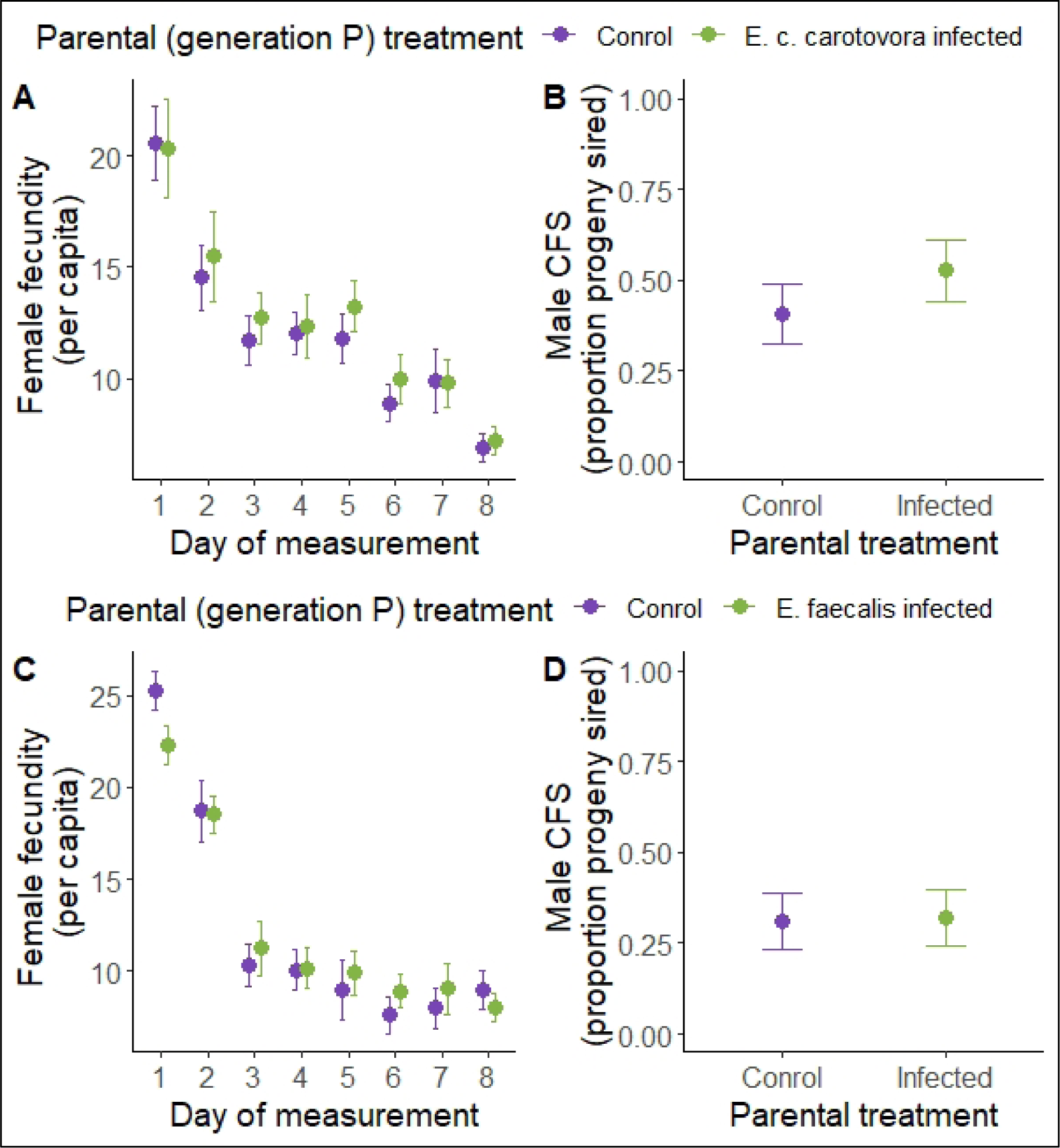
Reproductive output of first-offspring (F1) generation flies of infected and control parents in absence of an infection challenge. **(A)** Early-life fecundity of F1 females of *Erwinia c. carotovora* infected parents. **(B)** Competitive fertilization success of F1 males of *E. c. carotovora* infected parents. **(C)** Early-life fecundity of F1 females of *Enterococcus faecalis* infected parents. **(D)** Competitive fertilization success of F1 males of *E. faecalis* infected parents. Data pooled from both replicates for each pathogen. Y-axis error bars represent 95% confidence intervals around respective means.

Flies of parental generation (generation P) were infected with *E. faecalis*, and the reproductive output of F1 offspring was measured. Day of fecundity measurement had a significant effect on per capita female fecundity (F_1,318_ = 428.8211, p-value < 2.2e-16), but parental infection treatment did not have a significant effect (F_1,318_ = 1.4913, p-value = 0.223), and neither did the interaction between age × parental treatment (F_1,320_ = 1.9333, p-value = 0.165) (**figure 3.C**). Parental infection treatment did not have a significant effect on F1 male CFS (F_1,190_ = 0.0181, p-value = 0.893) (**figure 3.D**).

### 3.4. Systemic pathogen growth in first offspring (F1) generation flies after homologous infection challenge

Flies of parental generation (generation P) were infected with *E. faecalis* (*Ef*), and systemic pathogen load was measured for F1 offspring flies infected with the same pathogen. Systemic pathogen loads were measured for both infected flies (at 4- and 10-hours post-infection (HPI)) and sham-infected flies (at 4 hours post-infection). None of the sham-infected flies yielded any colony forming units.

In infected F1 females, parental infection treatment (F_1,192_ = 20.9157, p-value = 8.59 e-06), HPI (F_1,192_ = 150.7678, p-value < 2.2 e-16), and their interaction (F_1,192_ = 7.4957, p-value = 0.0068) had a significant effect on systemic pathogen load (**figure 4.A**). At 4 HPI, female offspring of *Ef*-infected parents (least-square mean, 95% confidence interval: 8.62, 8.01-9.23) carried similar bacterial loads as that of female offspring of control parents (LS mean, 95% CI: 9.17, 8.56-9.78). At 10 HPI, female offspring of *Ef*-infected parents (LS mean, 95% CI: 11.48, 10.87-12.09) carried a significant lower bacterial load compared to female offspring of control parents (LS mean, 95% CI: 13.67, 13.06-14.28).

**Figure 4.**
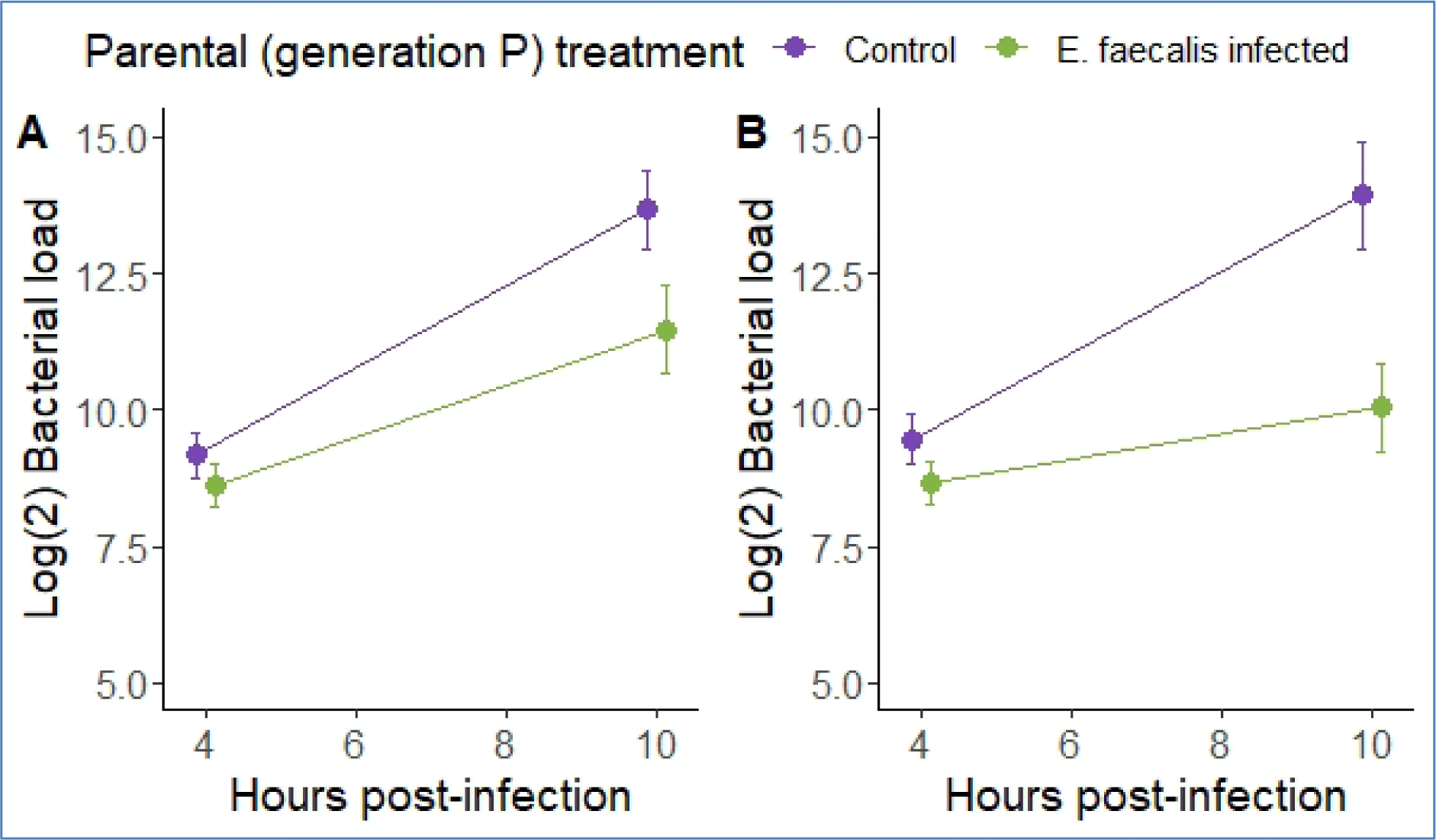
Systemic pathogen load in first-offspring (F1) generation flies of infected and control parents after a homologous infection challenge. Female **(A)** and male **(B)** offspring of *Enterococcus faecalis* infected parents were infected with *E. faecalis* and systemic bacterial loads were measured at 4- and 10-hours post-infection. Data pooled from all four replicates. Y-axis error bars represent 95% confidence intervals around respective means.

In infected F1 males, parental infection treatment (F_1,4_ = 39.649, p-value = 0.0032), HPI (F_1,4_ = 28.290, p-value = 0.0060), and their interaction (F_1,180_ = 23.079, p-value = 3.263 e-06) had a significant effect on systemic pathogen load (**figure 4.B**). At 4 HPI, male offspring of *Ef*-infected parents (LS mean, 95% CI: 8.65, 7.30-10.0) carried similar bacterial loads as that of male offspring of control parents (LS mean, 95% CI: 9.46, 8.11-10.8). At 10 HPI, male offspring of *Ef*-infected parents (LS mean, 95% CI: 10.05, 8.70-11.4) carried a significant lower bacterial load compared to male offspring of control parents (LS mean, 95% CI: 13.94, 12.59-15.3).

### 3.5. Post-infection fecundity of first offspring (F1) generation flies after homologous infection challenge

Flies of parental generation (generation P) were infected with *E. faecalis* (*Ef*), and post-infection fecundity of F1 female offspring was measured after infection with the same pathogen. When infected with *Ef*, female offspring of *Ef*-infected parents exhibited reduced mortality compared to female offspring of control parents (HR, 95% CI: 0.612, 0.499-0.751) (**figure 5.A**). No death was recorded in sham-infected F1 females from either parental treatments. Offspring infection status (F_1,4_ = 97.9970, p-value = 0.0006) had a significant effect on female fecundity, with infected females producing less progeny compared to sham-infected females (**figure 5.B**). Parental infection treatment (F_1,938_ = 0.6072, p-value = 0.4360) and interaction between parental and offspring infection treatments (F_1,938_ = 1.9811, p-value = 0.1596) had no effect on female fecundity (**figure 5.B**).

**Figure 5.**
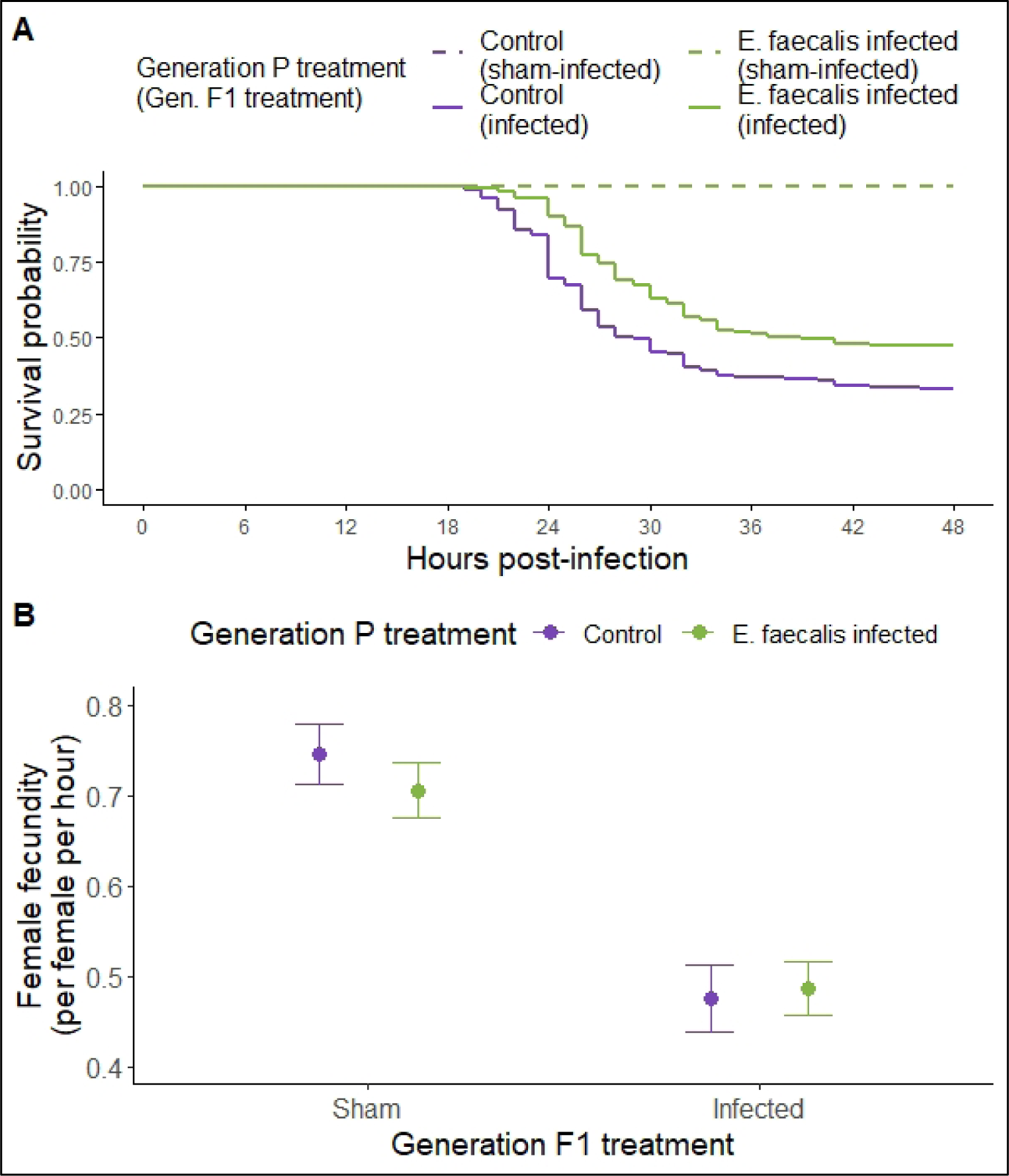
Post-infection fecundity of first-offspring (F1) generation flies of infected and control parents after a homologous infection challenge. Female offspring of *Enterococcus faecalis* infected parents were infected with *E. faecalis* (or subjected to sham-infections), and **(A)** post-infection survival and **(B)** post-infection fecundity of individual females were measured. Survival curves plotted using Kaplan-Meier method after pooling data from all four replicates. Data pooled from all four replicates for fecundity plot. Y-axis error bars represent 95% confidence intervals around respective means.

## 4. Discussion

### 4.1. Parental infection leads to increased offspring survival following a homologous infection challenge

We infected *Drosophila melanogaster* flies with two bacterial pathogens, *Erwinia c. carotovora* and *Enterococcus faecalis*, both of which are known natural pathogens of this species (Troha and Buchon 2009) and are therefore suitable for testing parental effects (Roth et al., 2009). For both pathogens, we found that first-generation (F1) offspring of infected parents were better at surviving an infection challenge, compared to the offspring of control parents, when they are infected with the same pathogen used to infect the parents, i.e., a homologous infection challenge. Improvement in offspring post-infection survival was observed for both sexes (**figure 1.A and 1.C**). There are two noteworthy nuances associated with these observations. *One*, improvement in offspring post-infection survival was only observed for offspring produced close to the point of infection in the parental generation. And *two*, the beneficial effect of parental infection on offspring post-infection survival was absent in the second-generation (F2) offspring.

For parents infected with *E. faecalis*, offspring produced between 48 and 66 hours post-infection had better post-infection survival compared to control offspring (**figure 1.C**), while offspring produced between 96 and 114 hours post-infection did not show similar improvement in post-infection survival (**figure 1.E**). Previous studies have demonstrated that the effect of parental infection on offspring immune function and life-history can depend upon the time of offspring production relative to the point of infection in *Tenebrio molitor* (Zanchi et al., 2011, Zanchi et al., 2012) and *Tribolium sp.* (Tate and Graham 2015). The fact that beneficial effects of parental infection on offspring traits are only observed in case of broods produced closer to the point of infection might potentially suggest that transmitting information regarding parental infection to the offspring is costly to the parents. Alternatively, it may suggest that infection induced changes in parents’ physiology are alleviated with time or the levels of pathogen associated molecular patterns in parents’ body decay with time, and therefore offspring produced at later time points do not inherit the information about parental infection. Which of these is true will depend upon the exact mechanism of transmission of parental effects to the offspring (Roth et al., 2018, Tetreau et al., 2019), which is yet to be completely elucidated (Contreras-Garduno et al., 2016, Prakash and Khan 2022).

Irrespective of if the parents were infected with *E. c. carotovora* or *E. faecalis*, the F2 offspring of infected parents did not differ in post-infection survival from the F2 offspring of control parents, barring one exception: F2 females of parents infected with *E. faecalis*, which exhibited improved post-infection survival (**figures 1.B, 1.D, and 1.F**). This suggests that in most scenarios the beneficial effect of parental infection on offspring survival persists only for one generation. For plastic parental effects to influence the evolution of any organismal trait, these effects need to be stably inherited across multiple offspring generations (Charlesworth et al., 2017, Yin et al., 2019). Since in our experiments we find that parental effects of pathogenic infections are stable for only one generation, parental effects are unlikely to influence the evolution of immune traits in *D. melanogaster* in a canonical, genetic assimilation based, plasticity-first mode of evolution (Bonduriansky and Day 2009). Alternatively, we hypothesize that parental effects in this species might prevent host population extinction in face of an emerging pathogen and help maintain sufficient population size for the hosts to have an opportunity to adapt to the invading pathogen (Bonduriansky and Day 2009). This has been previously suggested by some theoretical studies (Tate and Rudolf 2011, Tidbury et al., 2012) and is known to happen in the case of mistranslation-induced phenotypic plasticity in bacteria (Samhita et al., 2020).

A potential drawback of our experimental design is that since we subjected parental generation flies to pathogenic infections, with some amount of lethality, and collected eggs from these parents after the acute mortality phase (**supplementary figure S1**), we may have by default imposed selection for genotypes better at surviving pathogen challenges. Biased representation of such genotypes in the offspring generation due to survivor bias in the parental generation can also lead to increased post-infection survival in the offspring generation without any plastic parental effects. The two observations discussed above, in addition to their individual importance, together suggest that the increased post-infection survival in the offspring generation observed in our experiments are not driven by selection bias in the parental generation. Because, if the improvement in offspring survival was due to enrichment of *more immune* genotypes amongst the offspring, the observed improvement would not have been dependent on the time of egg collection (because at both 48 and 96 hours post-infection only the surviving parents contribute eggs) and would have also been observed in F2 flies (since a genetic change would be stable across generations). Therefore, we are convinced that the observed improvement of post-infection survival in the offspring generation is a plastic change in phenotype, brought about by the virtue of parental exposure to pathogenic infections, phenomenologically similar to trans-generational immune priming (Little and Kraaijeveld 2004, Krutz 2005, Rutkowski et al., 2023).

Two previous studies that tested for the effects of parental infection on offspring immune function discovered that exposing female flies to bacterial infections, which are pathogenic to varying degrees, has no effect on offspring post-infection survival and capacity to limit systemic pathogen proliferation following a homologous infection challenge (Linder and Promislow 2009, Radhika and Lazzaro 2023). We find the opposite results, both for post-infection survival and for limiting systemic pathogen proliferation (**see section 4.2**) in the F1 generation in our study, and interestingly while using a pathogen common with one of the previous studies (Radhika and Lazzaro 2023 used *E. faecalis*) and using the same route of infection (i.e., systemic infection via septic injury). There are two differences between our study and both the previous ones which might be of consequence. *One*, only female parents were infected in the previous studies while we infected parents of both sexes. *Two*, previous studies used inbred fly lines for their experiments while we obtained our experimental hosts from an outbred (*albeit* lab adapted) fly population. Previous studies in insects have demonstrated that both these factors, sex of the infected parent (Zanchi et al., 2011) and host genetic background (Khan et al., 2016, Khan et al., 2019), can influence trans-generational immune priming. Based on the data at hand, we cannot confirm or negate if one or both of these factors contribute to the differences in experimental outcomes of the present and previous studies.

### 4.2. Parental infection leads to increased disease resistance in the offspring

Disease resistance (i.e., host ability to limit systemic pathogen proliferation) and tolerance (i.e., host ability to mitigate infection-induced somatic damage) together contribute towards increased host survival following a systemic infection (Raberg et al., 2009). Both these traits can be differentially affected by parental exposure to pathogenic infections, which can in turn have different epidemiological consequences (Paraskevopoulou et al., 2022, Prakash and Khan 2022).

Our results show that offspring of infected parents, after being exposed to a homologous infection challenge, are better at slowing down the increase in systemic pathogen load (**figure 4**). This increased resistance probably underlies the increased post-infection survival exhibited by offspring of infected parents. We cannot rule out the role of tolerance, though, since we did not directly test for change in disease tolerance in our experiments. Therefore, our results show that, in *D. melanogaster*, parental experience with bacterial infections increases disease resistance in the offspring. Since increased disease resistance in hosts can drive evolution of more virulent pathogens (Raberg and Stjernman 2012), due to its negative effects on pathogen prevalence, our results also suggest that parental effects of pathogenic infection can promote evolution of more virulent pathogens.

We have thus demonstrated that parental exposure to pathogenic bacterial infections can both improve offspring post-infection survival (**see section 4.1**) and disease resistance in *D. melanogaster*, both of which have been suggested as defining features of trans-generational immune priming by previous studies (viz., survival, Roth et al., 2010; disease resistance, Moret 2006). One must although note that our observation of increased resistance may only be applicable for bacterial pathogens, since in the same host species, trans-generational immune priming can lead to increase in offspring resistance (Ben-Ami et al., 2020) or tolerance (Paraskevopoulou et al., 2022) depending on the type of pathogen the parents are exposed to.

### 4.3. Parental infection can lead to increased offspring survival following heterologous infection challenges

Subjecting offspring generation flies to heterologous infection challenges serves two goals: one is to test for the specificity of the parental effect, and the other is to test for associated costs (see section 4.4). Specificity relates to the capability of the host defense responses to differentiate between pathogens. Specificity in the context of priming in invertebrate hosts has been a matter of debate for long (Kurtz 2005, Schmid-Hempel 2005, Hauton and Smith 2007), an issue that is still not fully resolved (Contreras-Garduno et al., 2016, Tetreau et al., 2019, Prakash and Khan 2022). Studies in insects have shown that priming across generations can be pathogen specific (viz., Roth et al., 2010), or at times can be completely non-specific (viz. Dhinaut et al., 2018a). Studies in *Daphnia* have additionally demonstrated that empirical observation of specificity of priming in the same host species can depend on both on the pathogen used to prime the parental generation and on the offspring trait being measured (Little et al., 2003, Ben-Ami et al., 2020).

In our study we find that infecting parents with pathogenic bacterial infections improves offspring survival with some amount of cross-reactivity (*sensu* Kurtz 2005), i.e., offspring of infected parents are better at surviving heterologous infection challenges, with the breadth of cross-reactivity being determined by the identity of the pathogen used to infect the parents (**figure 2**). Additionally, the observed cross-reactivity is not symmetric: offspring of *E. faecalis* infected parents were better at surviving a challenge with *E. c. carotovora* (**figure 2.E**), but the *vice versa* was not always true and was dependent on the offspring sex (**figure 2.B**). Overall, these results suggest that increased offspring survival in *D. melanogaster* against bacterial infections by virtue of parental exposure to pathogenic infections is neither entirely specific nor entirely generic.

The fact that improved offspring survival by virtue of parental effects exhibits some degree of cross-reactivity has multiple evolutionary and ecological implications. *One*, trans-generational immune priming is hypothesized to evolve only when parents and offspring are likely to encounter the same pathogen (Little and Kraaijeveld 2004, Kurtz 2005), but if parental effects are cross-reactive then this stringent prerequisite for evolution of immune priming is relaxed. *Two*, trans-generational immune priming is hypothesized to not be beneficial for migrating offspring, since these offspring are expected to encounter novel pathogens (Pigeault et al., 2016), but with cross-reactive parental effects, even migratory offspring can reap the benefits of parental infection experience. *Three*, one of the ecological benefits of trans-generational immune priming is that it can prevent pathogen spread in host populations (Tate and Rudolf 2011, Tidbury et al., 2012), and with cross-reactivity, this benefit can extend to novel, non-endemic pathogens.

Sexual dimorphism in susceptibility to diseases is another issue under active exploration, in insects in general (Kelly et al., 2018), and more specifically in *D. melanogaster* (Belmonte et al., 2020). Despite years of research, and much hypothesizing (viz., Rolff 2002, Stoehr and Kokko 2006, Nunn et al., 2009), we do not yet fully understand what all factors can drive sexual dimorphism in disease susceptibility. We observed in our experiments that parental exposure to pathogenic infections can alter offspring susceptibility to heterologous infections in a sex-specific manner.

For example, the male offspring of *E. c. carotovora* infected parents were better at surviving infections with *E. faecalis* and *P. entomophila*, but such decreased susceptibility to infections was not apparent in case of the female offspring. We observe a similar scenario in F2 offspring of *E. faecalis* infected parents subjected to a homologous infection challenge: the female offspring exhibit increased post-infection survival compared to the control offspring, while the male offspring do not. Therefore, our results suggest that in *D. melanogaster*, prenatal exposure to pathogenic infections can lead to qualitatively sexually dimorphic improvement in offspring disease susceptibility.

### 4.4. Offspring flies incur no reproductive costs due to parental effects

Increased disease resistance in the offspring by virtue of parental exposure to infections can impose costs on the offspring that can compromise individual fitness, alter host population dynamics, and limit the evolution of transgenerational immune priming (Little and Kraaijeveld 2004, Tilbury et al., 2012, Roth et al., 2018). These costs, in general, are expected to manifest in two forms. *One*, in offspring of infected parents, improved disease resistance can trade-off with offspring reproduction and other life-history traits (viz., Roth et al., 2010, Dhinaut et al., 2018a) due to energy and resource costs of immune system upregulation. *Two*, offspring of infected parents, by virtue of being resistant to the pathogen encountered by their parents, can be more susceptible to other pathogens (viz., Sadd and Schmid-Hempel 2009) due to trade-offs between different components of the offspring’s immune system. Parental infection can occasionally even increase offspring susceptibility to the very pathogen encountered by the parents (viz., Vantaux et al., 2014, Ben-Ami et al., 2020). Additionally, costs in the offspring generation are more likely to be detected when there is a mismatch between offspring and parent environments (Tetreau et al., 2019), that is either in absence of infection or in case of a heterologous infection challenge.

We observed in our study that the measured reproductive potential of offspring of infected parents did not differ from that of control parents, in case of both pathogens used, and for both female and male offspring (**figure 3**). This suggests that in *D. melanogaster*, offspring incur no reproductive costs of increased disease resistance by virtue of parents being subjected to pathogenic bacterial infection. We also found that offspring of infected parents do not become more susceptible to heterologous infections (**figure 2**), within the range of pathogens tested, suggesting an absence of costs in the form of increased susceptibility to novel pathogens. Therefore, it can be expected that evolution of trans-generational immune priming should not be limited by concomitant physiological costs in this species, although ecological costs (such as increased host resistance selecting for increased pathogen virulence; Raberg and Stjernman 2012) can still act to slow down evolution of trans-generational immune priming.

The observation that parental infection status in general has no effect on offspring reproductive potential serves another purpose beyond suggesting an absence of costs. Previous studies have suggested that trans-generational improvement of offspring disease resistance can be driven by non-immunological mechanisms (Roth et al., 2018), such as differential nutritional provisioning of offspring by infected and uninfected parents (Pigeault et al., 2016), which improves overall offspring quality and not specifically only immunocompetence. If that is the case, it should also lead to a general improvement in offspring condition, which can potentially manifest as increased reproductive potential. Our observation therefore suggests that infecting parents with pathogenic bacteria does not improve general offspring quality, suggesting an immune function specific effect of parental infection.

### 4.5. Benefits of parental effects are limited to offspring post-infection survival

Negative effects of infection on host fitness extends to detrimental effects on host life-history traits, including host reproductive potential (Abbate et al., 2015, Nystrand and Dowling 2020). Trans-generational effects of prenatal infection can potentially mitigate the negative effects of pathogenic infection on host reproduction (Little et al., 2003, Paraskevopoulou et al., 2022). We tested for this possibility in our experiments and found that in case of infection with *E. faecalis*, female offspring of both infected and control parents exhibited similar reduction in fecundity when subjected to infection, while exhibiting significant difference in survival. This suggests that in *D. melanogaster* beneficial trans-generational effects of parental infection is limited only to offspring post-infection survival, and does not improve offspring fecundity tolerance (*sensu* Vale and Little 2012).

### 4.6. Conclusion

In this study we have demonstrated that, in *D. melanogaster*, parental exposure to pathogenic bacterial infections increases offspring post-infection survival and disease resistance following a homologous infection challenge. The survival benefit is primarily observed in the first offspring generation, and decays in the next generation, suggesting a plastic response instead of a genetic change driven by survivor bias in the parental generation. This also suggests low likelihood of genetic assimilation based, plasticity-first evolution immune traits in this species. Alternatively, parental effects probably help prevent population extinction at first encounter with pathogens thereby providing the host population an opportunity to adapt. Additionally, the survival benefit is restricted to offspring derived from eggs collected close to the infection challenge in the parental generation, further ruling out survivor bias in parents as cause of improved offspring survival. Increased disease resistance in offspring also suggests that parental effects can reduce pathogen prevalence and therefore increase selection for increased virulence amongst the pathogen population.

Parental exposure to infections also occasionally increases offspring survival following a heterologous infection challenge, with the breadth of cross-reactivity being dependent on the pathogen used for infecting the parents. This suggests that parental effects can prevent host population decline even when hosts face novel, non-endemic pathogens, such as during pathogen spillovers or host migration to a new environment. Importantly, improved post-infection survival in offspring by virtue of parental effects does not impose any concomitant cost on the offspring, either in the form of reduced reproductive output or in the form of increased susceptibility to heterologous infection challenges. Therefore, evolution of parental effects in *D. melanogaster*, with respect to immune traits, is unlikely to be constrained by physiological costs. Interestingly, although parental exposure to infections alleviates negative effects of infection on offspring survival, such ameliorative effects are not seen in context of infection induced reduction in offspring reproductive capacity.

In summary, therefore, we have demonstrated that pathogenic bacterial infections can induce trans-generational immune priming in *Drosophila melanogaster* flies without any measurable cost.

## Figures

**Supplementary figure S1.**
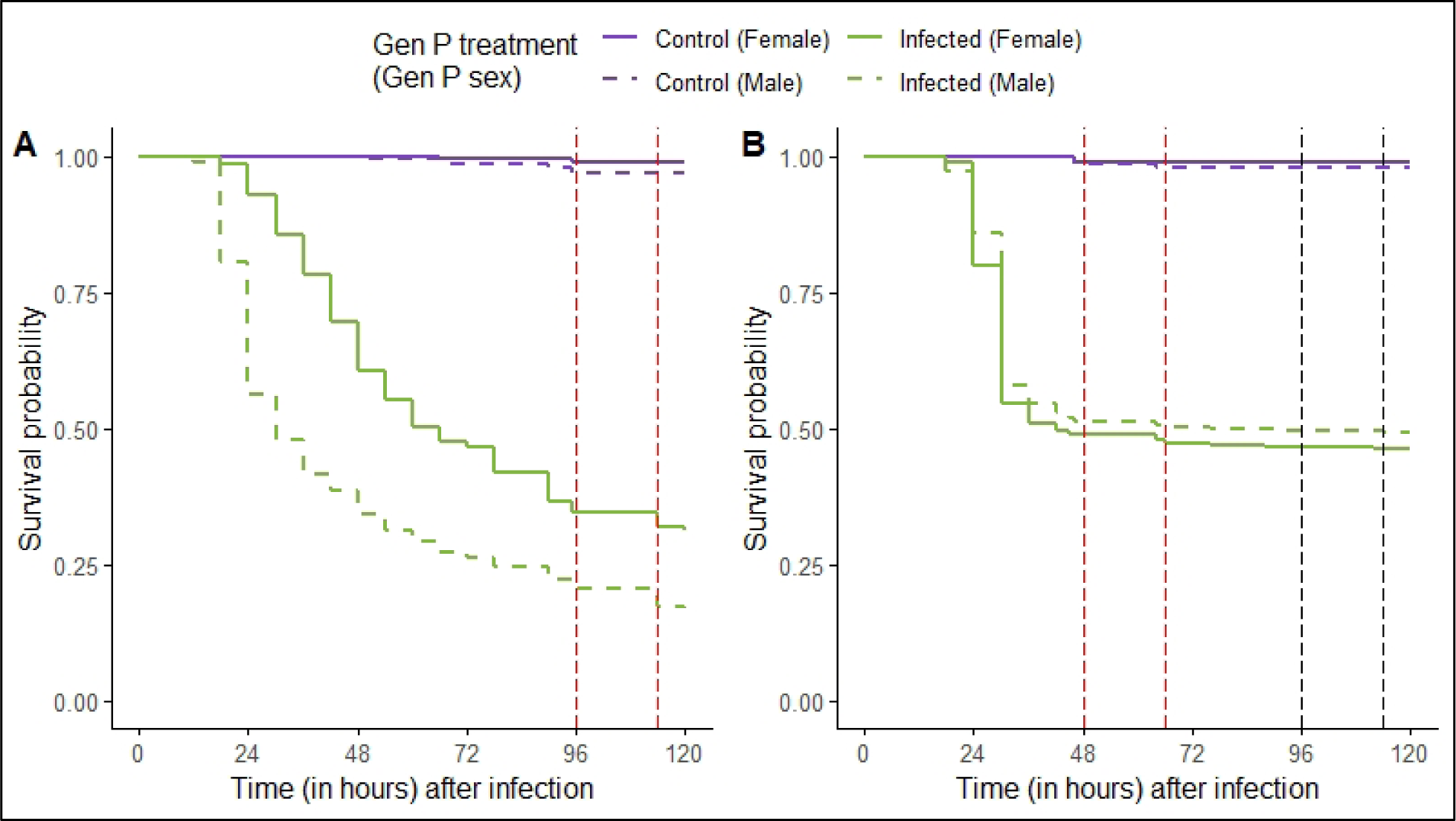
Representative plots for mortality of infected flies in the parental generation. **(A)** *Erwinia c. carotovora* and **(B)** *Enterococcus feacalis*. The egg collection window(s) are demarcated with dashed lines perpendicular to the X-axes.

**Supplementary figure S2.**
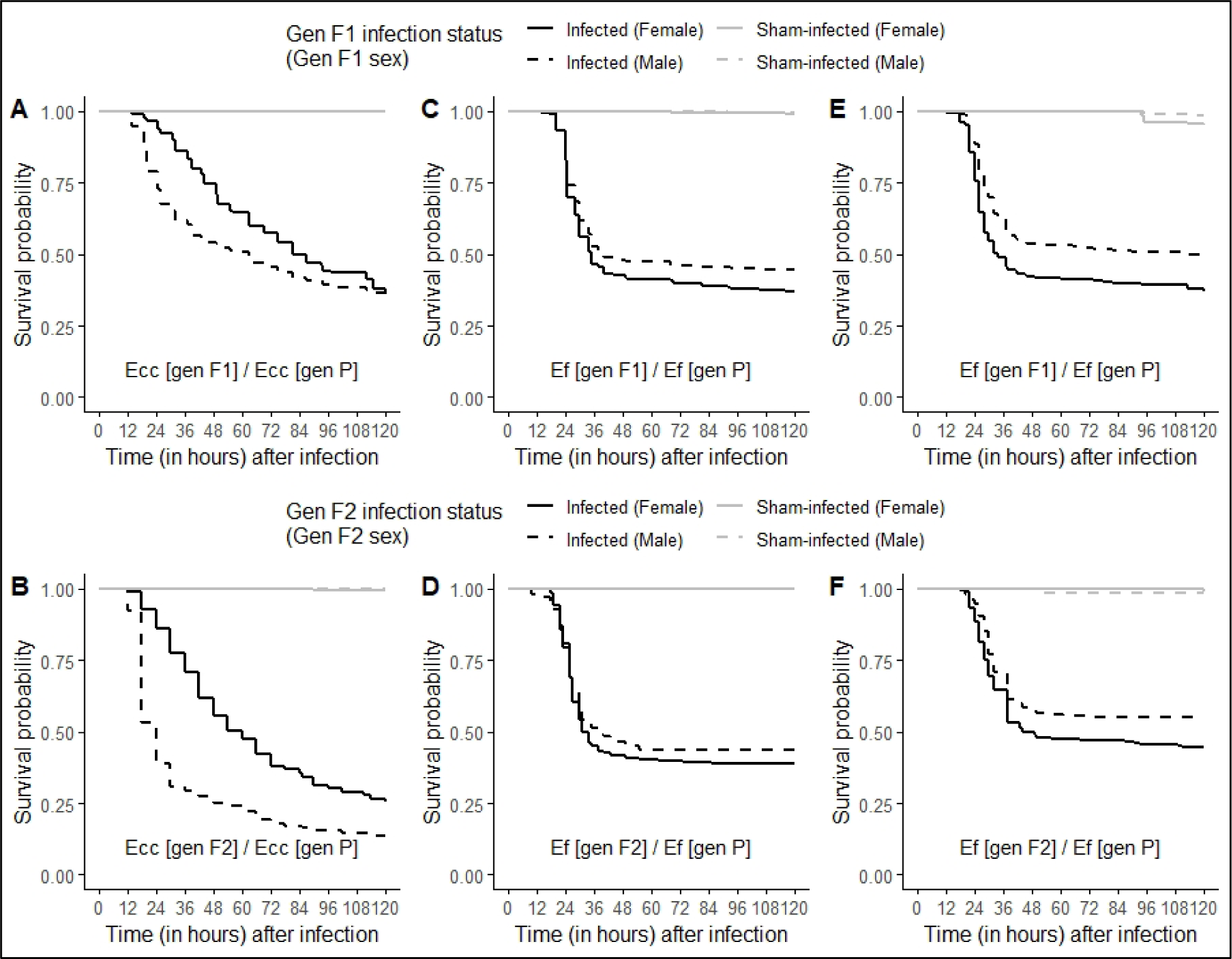
Plots for effect of homologous infection challenge on host survival in first (F1) and second (F2) offspring generations: comparison between sham-infected and infected flies with both parental treatments pooled together. [*Key: Pathogen used to infect F1 or F2 / Pathogen used to infect generation P.*]

**Supplementary figure S3.**
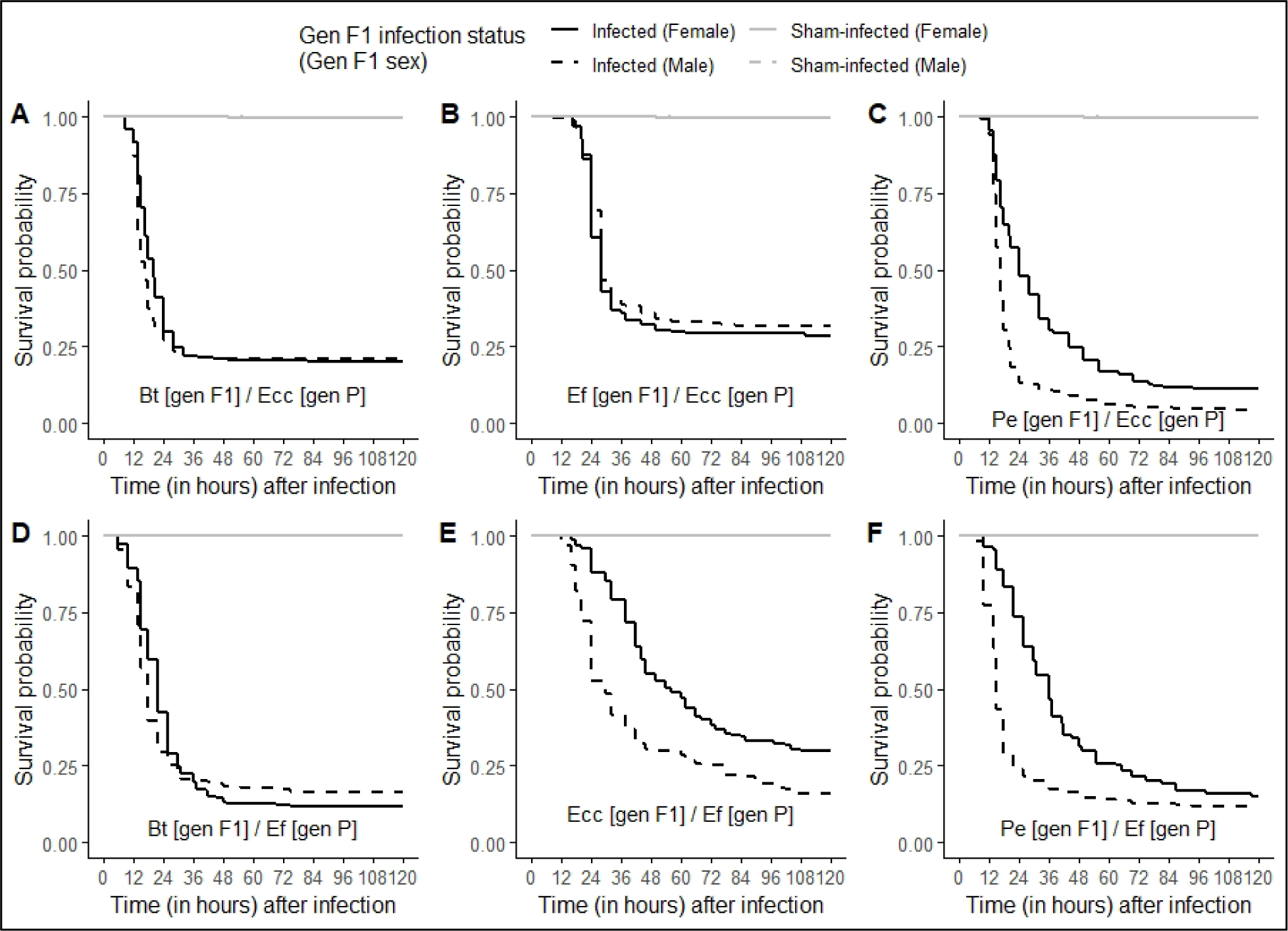
Plots for effect of heterologous infection challenge on host survival in first (F1) offspring generations: comparison between sham-infected and infected flies with both parental treatments pooled together. [*Key: Pathogen used to infect F1 / Pathogen used to infect generation P.*]

## Acknowledgement

The authors thank Dr. Aparajita Singh and Paresh Nath Das for logistic support during the execution of the experiments reported here, and Dr. Imroze Khan (Associate Professor, Ashoka University, India) for his comments on an earlier draft of the manuscript. The authors also thank Prof. P. Cornelis (Vrije Universiteit Brussel, Belgium) for providing the *Pseudomonas entomophila* isolate, Dr. Elio Sucena and Tania Paulo (Instituto Gulbenkian Ciencia, Portugal) for providing the *Erwinia c. carotovora* isolate, and Prof. Brian Lazzaro (Cornell University, USA) for providing the *Enterococcus faecalis* isolate, used in the experiments. The data presented in this manuscript was previously included as part of Aabeer Basu’s PhD thesis which can be accessed here (link: http://hdl.handle.net/123456789/5345).

